# A model Roseobacter employs a diffusible killing mechanism to eliminate competitors

**DOI:** 10.1101/766410

**Authors:** Garrett C. Sharpe, Scott M. Gifford, Alecia N. Septer

## Abstract

The roseobacter clade is a group of α-proteobacteria that have diverse metabolic and regulatory capabilities. They are abundant in marine environments and have a substantial role in marine ecology and biogeochemistry. However, interactions between roseobacters and other bacterioplankton have not been extensively explored. In this study, we identify a killing mechanism in the model Roseobacter *Ruegeria pomeroyi* DSS-3 by competing it against a group of phylogenetically diverse bacteria. The killing mechanism involves an unidentified antimicrobial compound that is produced when cells are grown on both surfaces and in suspension and is dependent on cell density. A screen of random transposon mutants revealed the killing phenotype, as well as resistance to the antimicrobial, require genes within an ~8 kb putative γ-butyrolactone synthesis gene cluster, which resembles similar pheromone-sensing systems in actinomycetes that regulate secondary metabolite production. Transcriptomics revealed the gene cluster is highly upregulated in wild-type DSS-3 compared to a non-killer mutant when grown in liquid coculture with a roseobacter target. Our findings show that *R. pomeroyi* has the capability to eliminate closely- and distantly-related competitors, providing a mechanism to alter the community structure and function in its native habitats.

## Introduction

Roseobacters are abundant members of marine microbial communities, comprising up to 20% of coastal and open ocean communities (1). Their prevalence in the marine environment is largely attributed to their diverse functional capabilities, including a wide range of metabolic strategies (2). Given their prevalence and metabolic versatility, roseobacters can have a substantial role in marine biogeochemical cycles, including the breakdown and release of carbon, sulfur, and other elements (3–10). Although studies have investigated the genes and regulatory mechanisms governing roseobacters’ biogeochemically important functions, less is known about how roseobacters interact with other bacteria to influence community structure and function (2, 4, 11–13). Specifically, the factors and mechanisms that may allow certain roseobacter strains to dominate a community remain largely unknown.

One strategy to increase fitness within a community is interference competition, in which a molecular mechanism is employed to kill or inhibit competitors. Some roseobacter strains have the capacity to produce antimicrobials and behave antagonistically toward other bacteria (14, 15). *Leisingera sp.* JC-1, isolated from the surface of Hawaiian bobtail squid egg clutches, produces the antimicrobial indigoidine, which differentially inhibited various *Vibrio* species and was hypothesized to protect the eggs from fouling microorganisms (16). *Phaeobacter gallaeciensis* produces the antimicrobial tropodithietic acid (TDA) when grown in coculture with the coccolithophore *Emiliana huxleyi*. TDA protects the algae from pathogens, and *P. gallaeciensis* receives nutrients in return. When the algal bloom begins to senesce, *P. gallaeciensis* produces anti-algal compounds known as roseobacticides that cause the mutualistic relationship to transition to parasitism (17). Finally, extracts from 14 roseobacter strains were found to produce compounds that inhibit the γ-proteobacterium *Vibrio anguillarium* (15). As these examples demonstrate, antimicrobial and anti-algal production by roseobacters can support beneficial relationships with their symbiotic partners and prevent competitor bacteria from gaining a foothold in preferred niches. Given that roseobacters are known to make up a large part of marine microbial assemblages and represent a mostly unexplored source of antimicrobials, further study of roseobacter interference competition mechanisms is important to understand how this diverse group shapes the communities they inhabit, which ultimately impacts the ecological services that these communities provide.

In this study, we used coculture assays to determine the potential for model Roseobacter *Ruegeria pomeroyi* DSS-3 to kill competitors and then employed a random mutagenesis approach to identify the killing mechanism. We chose *R. pomeroyi* because it has a relatively large genome (4.2 Mb) encoding multiple metabolic pathways and accessory functions for its generalist lifestyle (18–20). It is best known as a model organism for studying marine carbon and sulfur cycling (21–23). Although a previous study found *R. pomeroyi* can produce inhibitory compounds (15), the mechanisms and targets for *R. pomeroyi* killing have not been investigated. In this study, we describe a diffusible, cell density-dependent killing mechanism that DSS-3 uses to outcompete a phylogenetically diverse range of marine bacteria. The genes required for killing competitors appear to be horizontally acquired and are a fitness cost to DSS-3, suggesting that these genes may be selected for in the environment.

## Materials and Methods

### Growth of bacterial strains

Bacterial strains were grown on ½ YTSS plates supplemented with the appropriate antibiotic at 29°C, with the following two exceptions. *V. fischeri* ES114 pVSV208 was grown on LBS plates supplemented with the appropriate antibiotic at 24°C, and *E. coli* DH5α pGS001 was grown on LB plates supplemented with the appropriate antibiotic at 37°C. See Supplemental Information for additional experimental details including isolation of competitor strains, construction of tagged variants, and antibiotic concentrations used for selection.

### Surface Coincubation Assay

Individual colonies for each strain were placed into liquid media and grown while shaking at 200 rpm for 24 hours, and then subcultured and grown overnight in the same conditions. Overnight cultures were pelleted, and cells were resuspended in ½ YTSS media and diluted to an optical density at 600 nm (OD_600_) of 1.0. The two competing strains were mixed at a 1:1 or 9:1 DSS-3: target OD_600_ ratio, and 5 µL of the mixture was spotted on ½ YTSS plates and incubated (29°C, 24 hrs). The starting population of each strain was quantified by plating serial dilutions onto ½ YTSS plates supplemented with antibiotics selective for each strain, and colony forming units (CFUs) counted. After 24 hours, each coincubation spot was resuspended in 1 mL ½ YTSS medium and quantified by plating serial dilutions.

Filter separation coincubations were set up by spotting 20 µL of an OD_600_ 1.0 culture of either differentially-tagged DSS-3 or *Roseovarius sp.* TM1035, concentrated kanamycin antibiotic, or no addition (negative control) onto ½ YTSS plates, then placing a 0.2 µm nitrocellulose filter over the spot, and spotting 5 µL of tagged TM1035 (OD_600_=1.0) on top of the filter. The filter separation coincubations were then incubated at 29°C for 24 hours. The target strain was quantified at 0 and 24 hours by plating serial dilutions onto selective media as described above.

### Liquid suspension competitions

Strains were cultured overnight in 15 mL ½ YTSS liquid media containing the appropriate antibiotic at 29°C and shaken at 200 rpm. Cells were pelleted by centrifugation, the supernatant was removed, and the cell pellet was resuspended in 3 mL of fresh ½ YTSS. Each strain was then diluted to an OD_600_ of 0.2 in 10 mL ½ YTSS to achieve a 1:1 starting ratio. For the liquid competition assay testing the effect of cell starting densities, the 1:1 coincubation mixture was diluted with ½ YTSS medium to make 2-, 4-, 6-, 8-, and 10-fold diluted starting cocultures. Cell densities (CFU/ml) of each strain were then quantified by plating serial dilutions onto selective media at 0, 2, 4, 6, 8, and 24 hours.

### Transposon mutagenesis

The Tn5-Km transposon was conjugated into *R. pomeroyi* DSS-3 via coincubation with *E. coli* RH03 carrying the pUT mini Tn5-Km transposon delivery plasmid. Cultures were grown overnight in ½ YTSS at 29°C for DSS-3, and LB at 37°C supplemented with kanamycin and diaminopimelic acid (DAP), for *E. coli*. The two cultures were mixed at a 4:1 *E.coli* to DSS-3 ratio (vol/vol) and pelleted by centrifugation. The supernatant was removed, and the pellet was washed with ½ YTSS media and centrifuged again. All but 10 µL of the supernatant was removed, and this remaining supernatant was used to resuspend the cell pellet. The conjugation mixture was spotted on ½ YTSS DAP agar plates and incubated for 16 hours at 29°C. After 16 hours, the conjugation spot was resuspended in 1 mL ½ YTSS and centrifuged to pellet cells. The supernatant was removed, and the pellet was resuspended in ½ YTSS media and plated onto ½ YTSS agar plates supplemented with kanamycin. The plates were incubated at 29°C until kanamycin-resistant DSS-3 mutant colonies were visible on the plate (2-3 days incubation).

To screen for the non-killing mutant phenotype, each DSS-3 mutant was patched onto two corresponding sets of plates: a ½ YTSS plate supplemented with kanamycin and an agar overlay plate containing mCherry-tagged TM1035 target strain. Patches of DSS-3 mutants that retained the ability to kill the target generated a zone of clearing around themselves on target overlay plates, and were not examined further. Mutants that failed to create a zone of clearing were considered to be unable to kill, and those mutants were collected from the ½ YTSS kanamycin plate, cultured in ½ YTSS medium supplemented with kanamycin at 29°C while shaking at 200 rpm for 24 hours, and stored at −80°C for further characterization. Approximately 10 000 DSS-3 transposon mutants were generated and screened.

### inverse PCR (iPCR)

Mutants identified as losing the killing phenotype were cultured overnight in ½ YTSS media, and their DNA was extracted using the ZR Fungal/Bacterial DNA Miniprep Kit (Zymo Research, Ivrine, CA). The transposon insertion sites were then identified by iPCR (24). Briefly, 2 µg of mutant genomic DNA was digested overnight using the BssHII restriction enzyme (New England Biolabs, Ipswich, MA) in a 50 µL reaction and then cleaned and concentrated to 20 µL using the ZR DNA Clean and Concentrate-5 kit (Zymo Research, Irvine, CA). The resulting linear genomic DNA fragments were circularized using T4 DNA ligase (New England Biolabs, Ipswich, MA). The DSS-3 DNA sequence flanking the transposon was amplified with PCR by pairing a reverse primer that anneals at the 5’end of the transposon (5’-endseq, AS1193 or GS010) with a forward primer that anneals at the 3’ end of the transposon (3’-endseq, AS1196 or GS009). These primers anneal to the ends of the Tn5-Km transposon and amplify the circularized DNA fragment in between (see supplemental information for PCR cycles and conditions). Resulting PCR products were confirmed to be present using DNA electrophoresis, and the PCR products were cleaned and concentrated using the ZR DNA Clean and Concentrate-5 kit and Sanger sequenced by Eton Bio. The disrupted gene in each mutant was identified using NCBI BLAST. Mutation sites were confirmed by amplifying the transposon insertion site of uncut genomic DNA using a transposon-specific primer and a primer specific to the DSS-3 DNA near the mapper insertion site (See Supplemental).

### Transcriptomes

Liquid suspension competitions were set up as described above by mixing either DSS-3 Kn or DSS-3 SPOA0342 mutant (GCS64) with TM1035 pBBR1MCS-5 at a 1:1 ratio in triplicate, 10 mL cocultures (each strain at a starting OD_600_ of 0.2). The competition assay was subsampled regularly for 24 hours to determine strain population densities via serial dilutions on selective media plates. At 1.5 hours after starting the experiment, cells were collected by filtering 5 mL of coculture through a 0.22 µm polyethersulfone filter, and the filters were flash frozen in liquid nitrogen and stored at −80°C. RNA was extracted from the filters using the Mirvana RNA Extraction kit (Thermo Fisher Scientific, Walthon, MA). Residual DNA was removed using the TURBO DNA-free Kit (Invitrogen, Carlsbad, CA). cDNA libraries were prepared using the ScriptSEQ v2 kit and barcodes (Epicentre, Madison, WI) and sequenced with the HISEQ4000 platform (PE 50×50). Quality scores for each sequence were calculated using FastQC, and low quality sequences were removed from the raw sequencing data using Trimmomatic (sliding window trimming with average quality score lower than 20 across 4 bases removed). Reads were mapped to the DSS-3 genome using Bowtie 2 and the count intervals tool in Galaxy. Genes with statistically different relative abundances were identified with DESeq using the DE App deSEQ2 (25). Detailed parameters for the workflow are available in the Supplemental Information.

## Results

### *R. pomeroyi* kills a phylogenetically diverse range of bacteria

To determine whether *R. pomeroyi* exhibited antagonistic behavior toward other roseobacter strains, competitive assays were conducted in which DSS-3 was tested against six different roseobacter strains: *Sagittula stellata* E-37 (5), *Roseovarius sp.* TM1035 (26), *Sulfitobacter sp.* RAM1190, *Phaeobacter caeruleus* ANS2052, *Phaeobacter daeponensis*, and *Ruegeria sp.* RAM1602. DSS-3 was chromosomally tagged with a transposon expressing a Kanamycin-resistance gene (DSS-3 Kn) and mixed with a differentially tagged competitor strain. DSS-3 Kn and the competitor strains were initially mixed at a 1:1 ratio, based on optical density, and spotted onto ½ YTSS agar. The percent recovery of each strain was calculated by dividing the CFU count after 24 hours of coincubation by the initial CFU count at the start of the experiment. Recoveries >100% indicate the strain’s CFU count has increased over 24 hours, whereas <100% recovery indicates a strain has decreased. To determine whether a roseobacter strain is inhibited by DSS-3, the percent recovery of each strain with DSS-3 was compared to the recovery of the strain when incubated alone. After coincubation with DSS-3 at an initial 1:1 ratio, the recovery of every competing roseobacter was significantly lower when compared to its growth in monoculture, except for the control where DSS-3 was coincubated with itself, and for *P. daeponensis* (Student t-test, *P*<0.05, Fig. 1). However, when DSS-3 was mixed with *P. daeponensis* at a 9:1 ratio, this resulted in a similar reduction of *P. daeponensis* recoveries as observed with the other roseobacter strains (Fig. 1).

**Fig 1.**
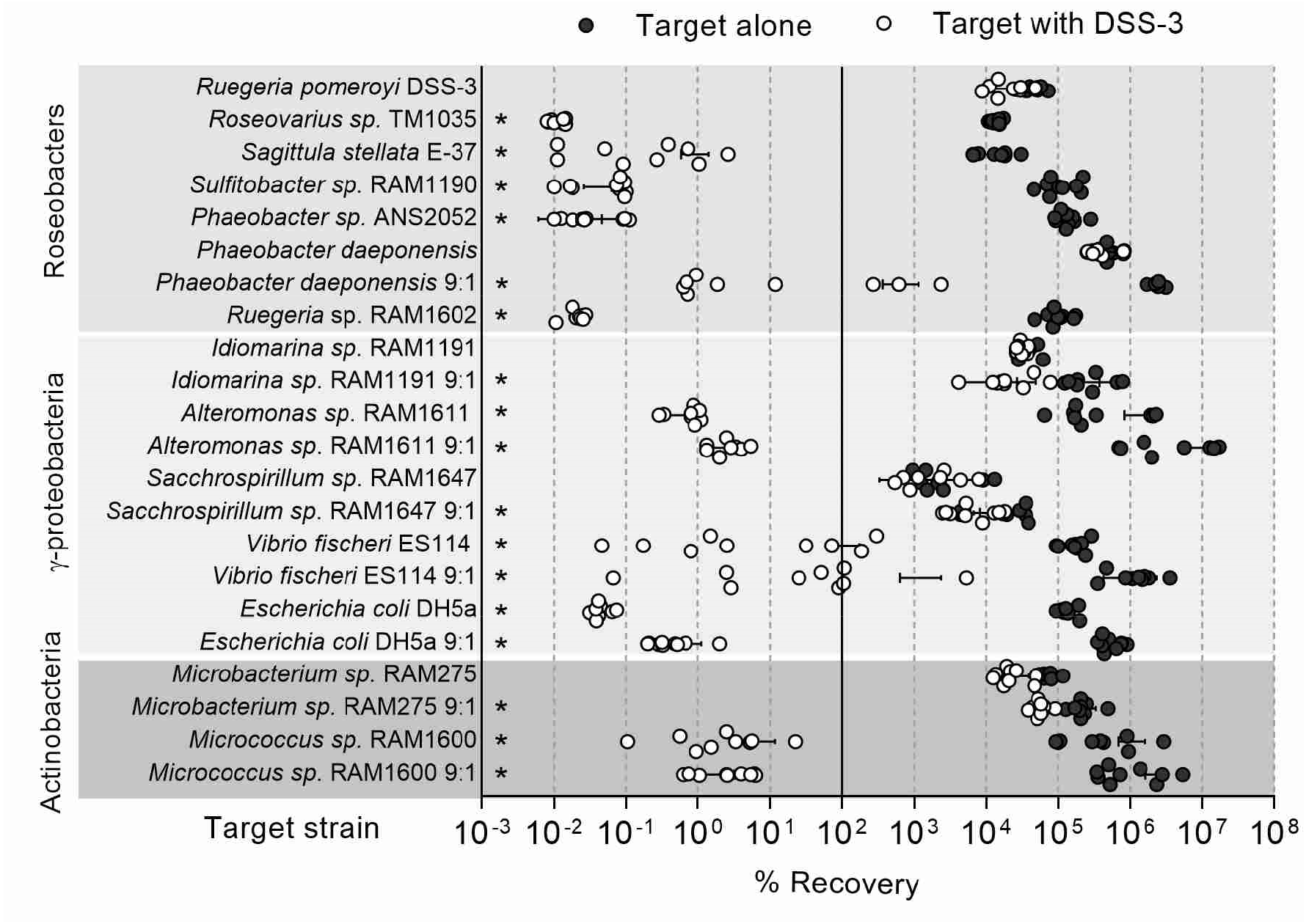
Effect of coincubation with *Ruegeria pomeroyi* DSS-3 on the growth of phylogenetically diverse bacteria. Percent recovery for each species after 24 hours when incubated in monoculture with itself (black) or in coculture with DSS-3 (white) on ½ YTSS agar, where less than 100% recovery corresponds to a decrease from the initial colony forming unit (CFU) count, and greater than 100% recovery corresponds to an increase from the initial CFU count. Asterisks denote a *P*<0.05 using a students t-test comparing percent recovery of each species alone to percent recovery when coincubated with DSS-3. Error bars indicate standard deviation.

To determine whether DSS-3 can inhibit more distantly related bacteria, several marine γ-proteobacteria and actinobacteria were selected as competitors: *Alteromonas sp*. RAM1611, *Saccharospirillium sp.* RAM1647, *Idiomarina sp*. RAM1191, *Vibrio fischeri* ES114, *Escherichia coli* DH5α, *Microbacterium phyllosphaerae* RAM275, and *Micrococcus sp.* RAM1600. These strains were initially mixed at a 1:1 ratio (DSS-3: competitor) as done with the roseobacter coculture assays, but 9:1 ratios were also performed to determine if inhibition occurs at higher starting DSS-3 cell densities. In the 1:1 ratio experiments, the *Idiomarina, Saccharospirillium*, and *Microbacterium* strains did not show statistically significant decreases in recovery when coincubated with DSS-3 Kn, whereas the *Alteromonas, Vibrio, Escherichia*, and *Micrococcus* strains did (Student t-test, *P*<0.05). In the 9:1 ratio experiments, all y-proteobacteria and actinobacteria strains tested had significantly lower recoveries when cocultured with DSS-3 Kn versus in monoculture (Fig. 1). Taken together, these data suggest that DSS-3 uses an unknown mechanism to kill competitor strains during growth on surfaces.

### *R. pomeroyi* kills competitors in suspension

To determine whether DSS-3 killing can also occur in liquid suspension, DSS-3 was coincubated for 24 hrs with the susceptible target strain *Roseovarius sp.* TM1035 in liquid medium at a 9:1 starting ratio (DSS-3:TM1035), and CFUs were quantified for each strain at various time points during the 24 h coincubation. Two hours after the start of the incubation, TM1035 CFUs were reduced by four orders of magnitude when coincubated with DSS-3 in liquid medium, and by 4 hours the TM1035 abundance was reduced below the limit of detection (200 CFUs/ml) (Fig. 2A).

**Figure 2.**
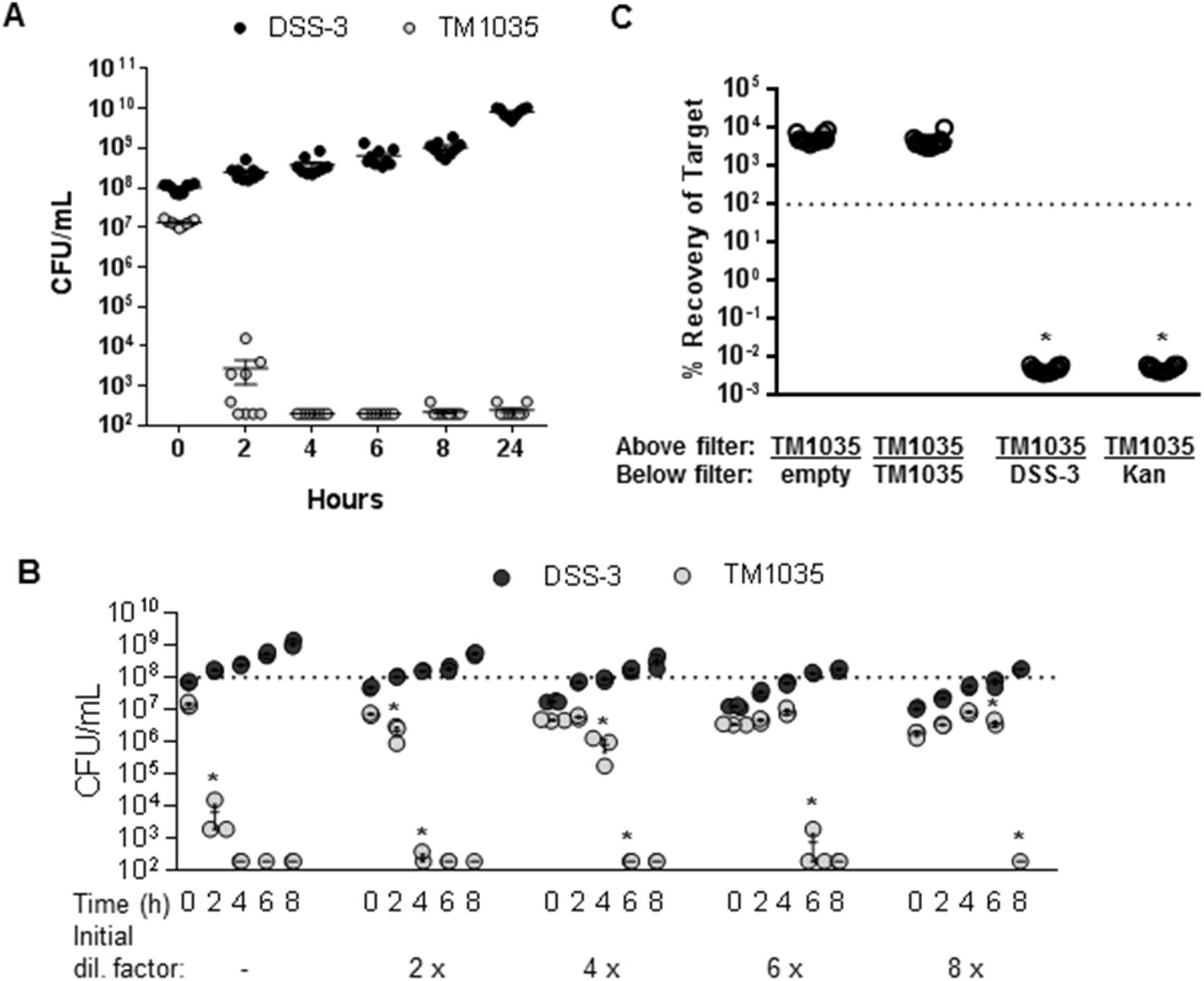
DSS-3 can kill in suspension using a diffusible, density-dependent mechanism. Cell density (CFUs/ml) of DSS-3 Kn (black) and TM1035 (gray) in liquid coculture. (A) All 9 replicates for the undiluted liquid competition assay, and (B) a representative experiment for liquid dilution experiment. Asterisks denote time points where TM1035 CFU/mL are statistically lower than that of the previous time point (*P*<0.05, Student’s t-test). (C) Percent recovery of tagged TM1035 at 24 hours when grown on a filter above: an empty control, a differentially tagged TM1035 strain, DSS-3, or 2 µg kanamycin antibiotic. Asterisks denote a *P*<0.0001 using a Student’s t-test comparing each experimental condition to the TM1035/empty control. Error bars indicate standard deviation, although some are too small to be seen.

Given that some killing mechanisms are dependent on cell density (27–34), and we sometimes only observed inhibition when DSS-3 initially outnumbered its competitor (Fig 1), we next examined whether the DSS-3 killing phenotype might be correlated with population density in culture. Liquid competition assays were conducted as above, except the starting densities of both DSS-3 and TM1035 in the coculture were diluted by 2-, 4-, 6-, and 8-fold. If diluting the initial coculture cell density delays DSS-3 killing, then killing activity requires a particular cell density to promote the killing function. For each dilution tested, a statistically significant reduction of TM1035 target cells did not occur until DSS-3 reached densities of ~10^8^ CFU mL^−1^ (Fig. 2B). This result suggests that, under the conditions used here, DSS-3 must achieve a cellular concentration threshold of >10^8^ cells mL^−1^ before killing is detected and that its antimicrobial function may be controlled in a density-dependent manner.

### *R. pomeroyi* uses a diffusible killing mechanism

Antimicrobials can function as diffusible molecules, as is the case for most conventional antibiotics, or they can require direct cell-cell contact for transfer from killer to target cells (35–38). To determine whether DSS-3’s killing phenotype is contact dependent or diffusible, DSS-3 was cocultured with TM1035 on agar plates as above, except the two strains were separated by a 0.22 µm nitrocellulose filter, which prevents physical contact between the two strains while allowing diffusible molecules to be exchanged. When tagged TM1035 was spotted on a filter with nothing below, or with untagged TM1035 below the filter, the percent recovery of tagged TM1035 was above 100% (Fig 2C). However, if tagged TM1035 was spotted onto a filter above DSS-3 or concentrated kanamycin antibiotic, TM1035 CFUs were reduced beyond the limit of detection (Fig. 2C). These results suggest that DSS-3 employs a diffusible killing mechanism that does not require direct contact with target cells.

### Random transposon mutagenesis yields non-killer DSS-3 mutants

To identify the genes and possible mechanism(s) required for DSS-3 killing, we generated a random transposon library and screened it for DSS-3 mutants that can no longer kill a competitor strain. The screen was based on the observation that when wild-type DSS-3 is grown on a lawn of fluorescently-tagged TM1035 target cells, a distinct zone of killing is observed around the DSS-3 colony (Fig 3A). If the transposon disrupts a gene required for killing, then the DSS-3 mutant colony will not produce a zone of killing. We screened 10 000 DSS-3 mutants and isolated seven non-killer mutants that were either unable to produce a zone of killing of TM1035, or displayed an intermediate zone of killing (Fig. 3A).

**Figure 3.**
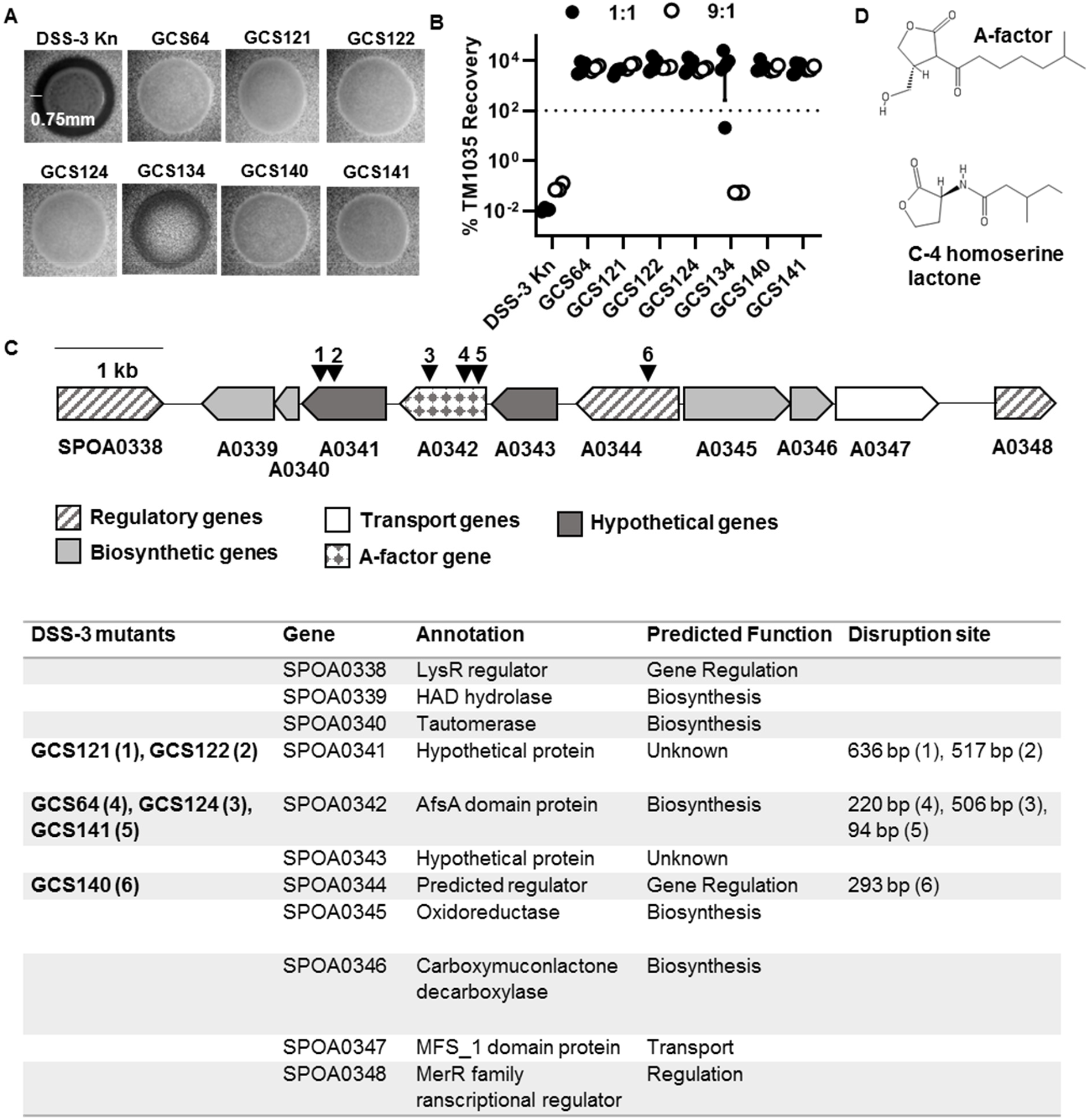
Characterization of DSS-3 mutants that have lost the killing phenotype. (A) Microscope images of DSS-3 Kn tagged wild-type and seven identified non-killer mutants when grown on fluorescent TM1035 agar overlay plates. Scale bar denotes length of inhibition zone if killing occurs. (B) Percent recovery of TM1035 after 24 hours of coincubation on surfaces with the DSS-3 Kn control or each potential non-killer mutant at a 1:1 (black circles) and 9:1 (white circles) killer to target strain OD_600_ ratio. Asterisks denote significant reduction (*P*<0.05, Student’t t-test) when compared to the DSS-3 Kn control. Error bars indicate standard deviation. (C) Diagram of putative GBL synthesis gene cluster on the megaplasmid. Black triangles and numbers correspond to transposon insertion mutants listed in the table below. (D) Structures for quorum molecules A-factor GBL and C-4 homoserine lactone.

To confirm these mutants could no longer kill, DSS-3 mutants were coincubated with TM1035 on agar surfaces, as described above using both a 1:1 and 9:1 starting ratio of DSS-3 to TM1035 target. Of the seven mutants, six were unable to kill TM1035 to a statistically significant degree after 24 hours at either starting ratio (Fig. 3B). However, one mutant (GCS134), which exhibited only partial killing on agar overlay plates (Fig. 3A), was not as efficient at killing target at a 1:1 starting ratio (Fig. 3B, filled circles), but was able to kill when initially outnumbering the target at a 9:1 starting ratio (Fig 3B, empty circles). These results suggest that six of the isolated mutants have lost the ability to kill, and one mutant (GCS134) displayed reduced killing ability that could be restored by increasing its population size at the beginning of the coincubation experiment.

### The genes required for killing are located in a putative γ-butyrolactone biosynthesis gene cluster

The locations of the transposon insertions were mapped using inverse PCR. For mutant GCS134, whose killing ability was restored at 9:1 starting ratio, the transposon insertion site mapped to a sulfate adynylyltransferase (ATP sulfurylase) gene (SPO0900), which is predicted to mediate the cellular assimilation of inorganic sulfur (39–41). We hypothesized that this mutant, which could not kill TM1035 cells at a 1:1 starting ratio (Fig 3B), has a reduced growth rate and could not achieve the threshold density required for killing. Indeed, the growth rate of GCS134 was reduced compared to the tagged DSS-3 Kn strain (Supplemental Fig 1A). Moreover, when the CFUs were calculated for both the mutant and TM1035 target strains in coculture, GCS134 was unable to reach the 10^8^ CFU ml^−1^ threshold when a 1:1 starting ratio was used, and did not eliminate TM1035 (Supplemental Fig 1B). However, when GSC134 was coincubated with TM1035 at a 9:1 starting ratio, the mutant could achieve the necessary cell density and eliminate TM1035 target after a 24 hour coincubation (Supplemental Fig. 1B). Because GCS134 retained its ability to kill when coincubation conditions permitted it to achieve a sufficiently high cell density, we did not consider this mutant in further analysis.

The six remaining mutants contained transposon insertions in a single gene cluster located on DSS-3’s megaplasmid (Fig. 3C). According to an antiSMASH analysis (42), this gene cluster encodes a potential γ-butyrolactone (GBL) biosynthesis gene cluster, including a predicted A-factor synthesis (AfsA) domain protein that is essential for GBL production in other bacteria (43, 44), additional biosynthetic proteins, three predicted transcriptional regulators, a predicted transporter, and two hypothetical proteins (Fig. 3C). GBLs, such as A-factor, are quorum sensing molecules with a similar structure to the well-studied quorum sensing molecule C-4 homoserine lactone (Fig. 3D) and have been found to regulate antibiotic production and cell differentiation in streptomycetes (45–49). Together, these data revealed two important findings: 1) they indicate our screen reached saturation because we obtained multiple, independent insertions in one gene cluster, with two genes having multiple, independent transposon insertions, and 2) the predicted GBL biosynthesis cluster is required for DSS-3 to kill a competitor strain.

### DSS-3 GBL homologs are found primarily in distantly related taxa

Given that GBL synthesis has primarily been described in Actinobacteria (46, 47), we examined whether DSS-3’s GBL gene cluster may have arisen through horizontal transfer from distantly-related bacteria. Homologs to the AfsA domain protein SPOA0342 and the flanking hypothetical proteins SPOA0341 and SPOA0342 were identified using BLASTx homology searches against NCBI’s non-redundant protein database in order to identify GBL presence in other bacterial clades (Supplemental Table S2). Of the 35 organisms found to encode a homolog of SP0A0342, 29 encode all three genes and 6 encode SPOA0342 and one flanking gene. Only five of these organisms are alphaproteobacteria, two of which are roseobacters (*Ruegeria sp.* EL01 and *Shimia marina*), and the majority of organisms that encode homologs to at least two of these three GBL genes belonged to the γ-proteobacteria and Actinobacteria (Fig. 4, Supplemental Table S2). We next searched more broadly for proteins containing the key A-factor domain (Pfam domain PF03756) using AnnoTree (50). Similar to the Blastx search results, the AnnoTree results revealed that ~93% of the total identified A-factor domain containing proteins were found in γ-proteobacteria and actinobacteria, and less than 2% were found in roseobacters (Fig. 4). Together, these results suggest that DSS-3’s GBL gene cluster may have been acquired horizontally from either a γ-proteobacteria or Actinobacteria lineage.

**Figure 4.**
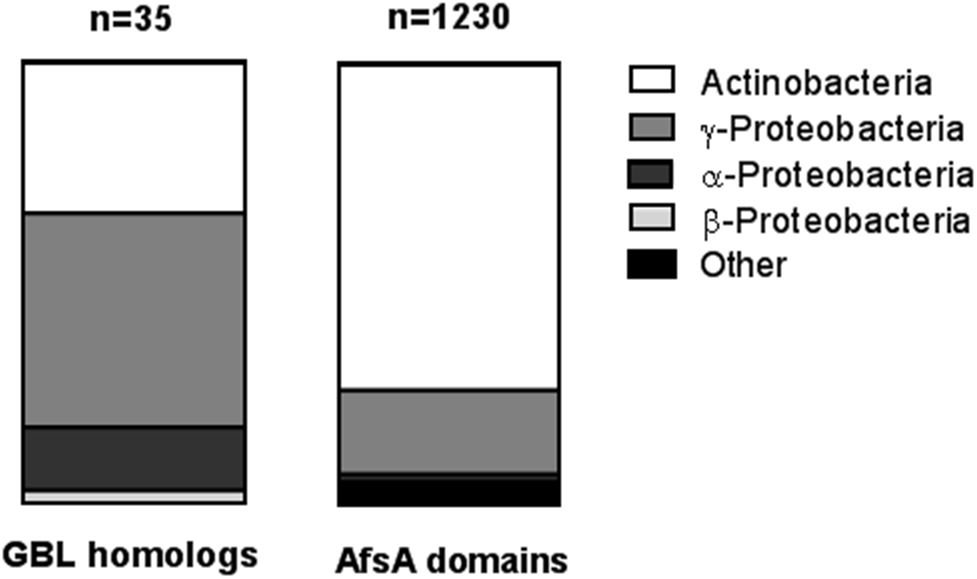
Presence of SPOA0341-0343 homologs and the AfsA domain (PF03756) in bacterial phyla. Homologs of genes SPOA0341, 0342, and 0343 with >30% identity were identified with Blastx using nr database (left). AfsA domain (PF03756) proteins were also identified using AnnoTree (right).

### GBL genes are required to protect against self-killing

Bacteria will often produce immunity factors to prevent self-killing while employing antimicrobials (51). Given that antimicrobial and immunity genes are often encoded near one another on the genome and coexpressed, our transposon insertions may have also disrupted DSS-3’s genetic factors for immunity. To determine whether the non-killer mutants had become sensitive to killing, we coincubated the mutant DSS-3 strains with a differentially-tagged parental DSS-3 strain using a 1:1 starting ratio and assayed percent recovery of each mutant after 24 hrs on surfaces. Although some mutants grew in the presence of the parent strain (>100% recovery), others were significantly inhibited or nearly eliminated. All three *afsA* (SPOA0342) mutants (GCS64, GCS124, and GCS141) showed no statistically significant reduction in percent recovery when coincubated with the parent strain, suggesting these strains retained immunity to the antimicrobial (Fig 5A). By contrast, both mutants with transposon insertions in the hypothetical protein SPOA0341 (GCS121 and GCS122) had reduced percent recoveries when coincubated with the wild-type (Fig. 5A), suggesting they are sensitive to antimicrobial production by the parent strain. Interestingly, we recovered ~100-fold less of SPOA0341 mutant GCS122 compared to SPOA0341 mutant GCS121, suggesting that although both mutants have a transposon insertion in the same gene, the insertion site and/or orientation of the transposon may also influence expression of immunity factors. Finally, the predicted regulator mutant (SPOA0344, GCS140) showed the highest sensitivity to killing, with the recovery of this strain reduced below the limit of detection after 24 hrs coincubation with the parent; a result similar to what we observed when coincubating other sensitive roseobacter isolates with differentially-tagged DSS-3. Taken together, these findings indicate that in addition to encoding the factors necessary for killing, the gene cluster also encodes genes required for immunity.

**Figure 5.**
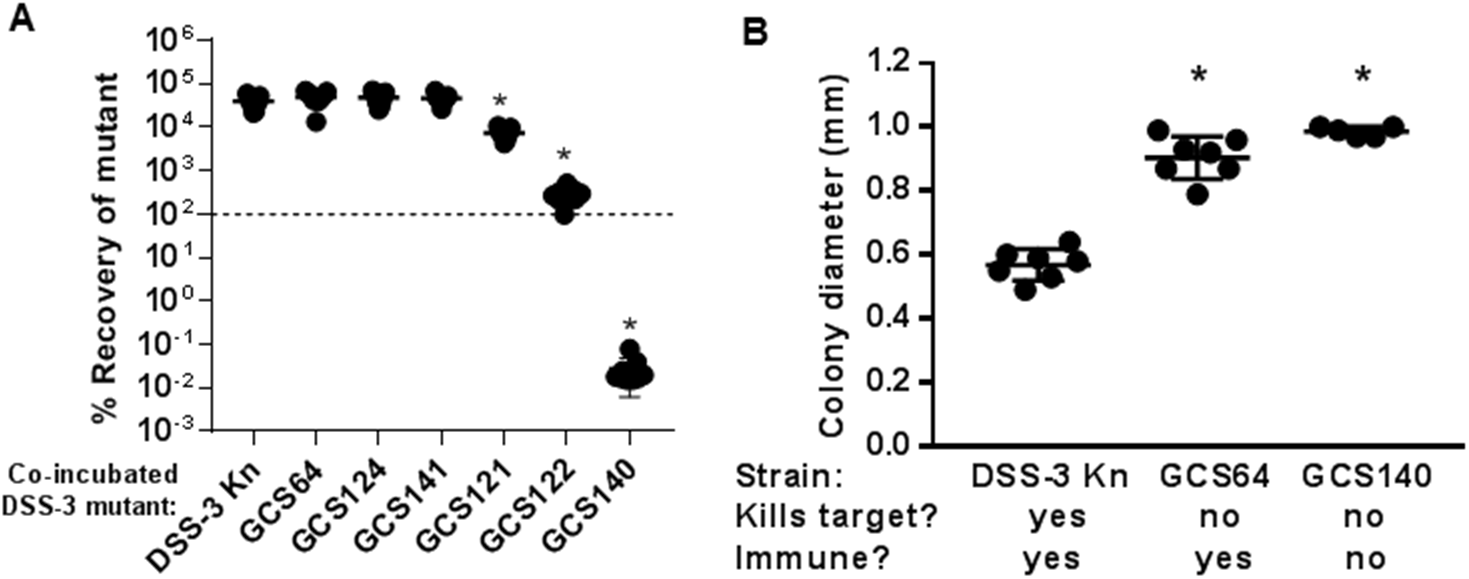
Killing ability, but not immunity function, is a fitness cost for DSS-3. (A) Percent recovery of DSS-3 Kn and each non-killer mutant when coincubated on ½ YTSS agar plates with differentially-tagged wild-type DSS-3 at a 1:1 DSS-3 wild-type to DSS-3 mutant ratio. Asterisks denote a *P*<0.001 using a Student’s t-test comparing mutant to wild type. (B) Colony diameter for DSS-3 Kn and select mutants grown on ½ YTSS agar plates for 48 hours. Asterisks denote a *P*<0.001 using a Student’s t-test when comparing each mutant to the wild-type. Error bars indicate standard deviation.

### The GBL gene cluster is energetically costly

In an analysis of the DSS-3 proteome, Christie-Oleza *et al.* (2012) found five proteins encoded in the GBL gene cluster (SPOA0339-0343) comprise 1-6% of the entire DSS-3 proteome under various conditions (52). Therefore, we hypothesized that the non-killer mutants would have a higher growth rate, either due to their reduced GBL protein synthesis or lack of antimicrobial production, which may also be energetically costly. To determine whether the mutants grew on surfaces more quickly than the wild-type, the DSS-3 Kn, SPOA0342 *afsA* mutant GCS64, and the SPOA0344 regulator mutant GCS140 were plated onto ½ YTSS agar and the diameters of the colonies for each strain were measured after 48 hours of incubation at 29°C. The colony diameter for two representative non-killer mutants (one that is immune and one that is no longer immune) were approximately twice that of the wild-type strain (Fig 5B). Taken together, these data suggest that killing ability, but not immunity function, may be a fitness cost for DSS-3.

### *afsA* is required for transcription of the GBL gene cluster

Because A-factor is known to regulate antimicrobial production in actinomycetes, we hypothesized that the *afsA* gene product may have a similar role in regulation of DSS-3 antimicrobial production. To determine the AfsA-dependent regulon in DSS-3, we compared the transcriptomes of the *afsA* mutant GCS64 and DSS-3 Kn when coincubated with *Roseovarius sp.* TM1035 in liquid suspension. The cocultured cells used for transcriptome sequencing were collected at 1.5 hours when killing of TM1035 by wild-type DSS-3 begins to occur in a 1:1 liquid coculture (Fig. 6AB). Of the 4 252 genes in the DSS-3 genome, only 21 genes were significantly differentially transcribed between DSS-3 Kn and the *afsA* mutant GCS64, 10 of which corresponded to the putative GBL cluster (Fig 6C). Transcripts of these ten genes were enriched by 4 to 60 fold in the wild-type strain compared to the *afsA* mutant. The only gene encoded in the GBL gene cluster that was not differentially expressed was the predicted regulator encoded in SPOA0344. Taken together, these data show 1) the *afsA* gene is required for expression of most of the genes in the GBL operon, and 2) this regulatory mechanism does not significantly impact expression of other genes in the DSS-3 genome.

**Figure 6.**
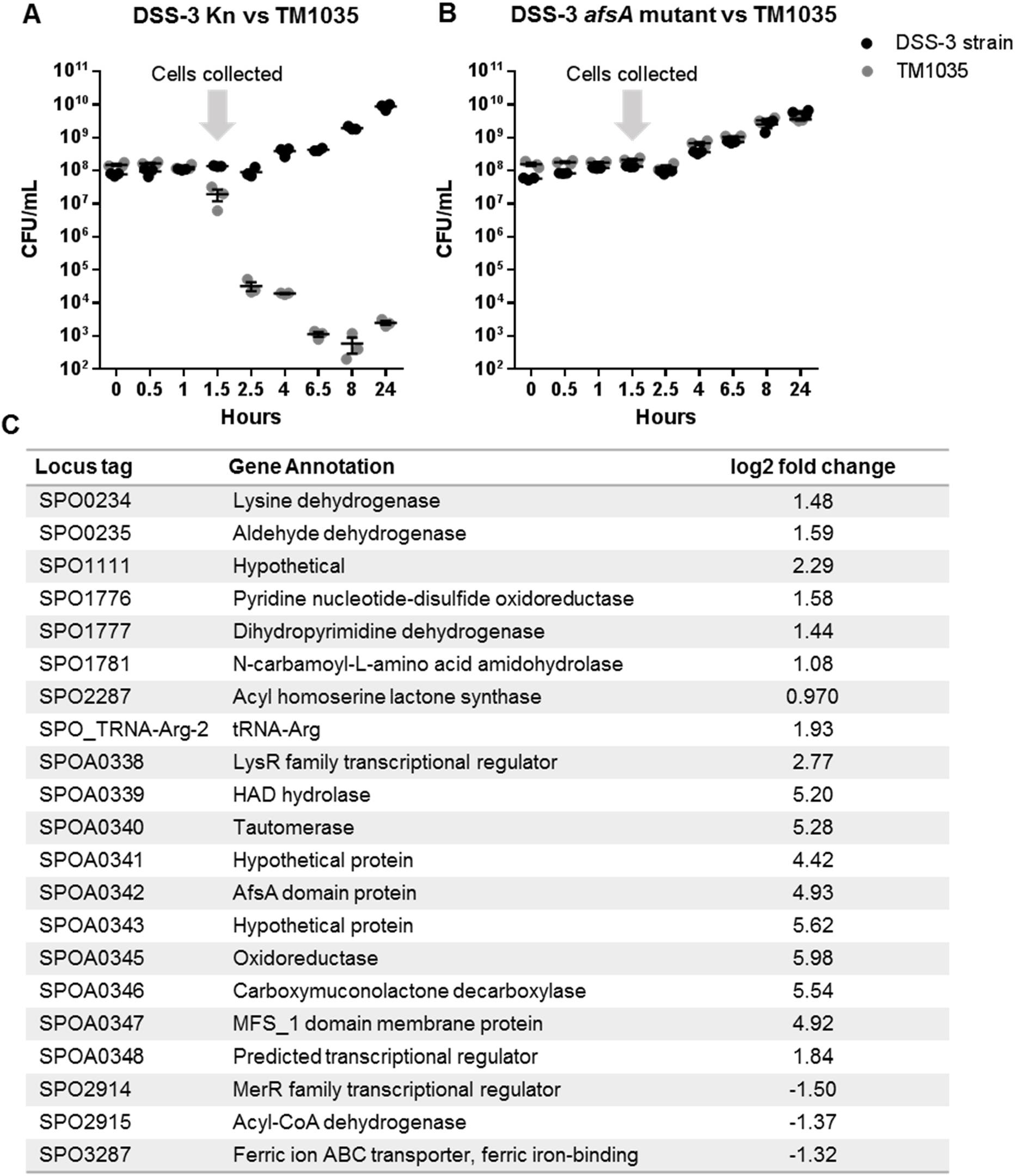
The *afsA* gene is required for transcription of GBL cluster genes in liquid coculture. (A and B) Collection of cells from coincubation experiments for transcriptome analysis of DSS-3 Kn and *afsA* mutant GCS64. DSS-3 Kn and DSS-3 mutant GCS64 were each co-incubated with target strain TM1035 at a 1:1 ratio (OD_600_ of 0.2 each) and quantified by serial dilution on selective media. Cells were collected on a 0.22µm polyethersulfone filter at 1.5 hours (arrow), and RNA was extracted from each filter and processed for transcriptome sequencing. (C) Statistically significant differentially regulated genes comparing the wild-type DSS-3 Kn and the SPOA0342 mutant GCS64 when coincubated with target Roseobacter strain TM1035. The log2 fold change denotes whether transcripts were enriched or depleted in the tagged wild-type strain DSS-3 Kn compared to the non-killer mutant GCS64.

### The GBL gene cluster is required for DSS-3 to outcompete phylogenetically diverse marine bacteria

To test the GBL gene cluster’s role in killing other bacterial types, DSS-3 Kn and the *afsA* mutant GCS64 were used in competition assays with the full taxonomic range of competitor strains described in Figure 1. For the roseobacter competitions, a 1:1 starting ratio was used for all the roseobacter competitions except for *P. daeponensis*, in which case a 9:1 DSS-3 to *P. daeponensis* ratio was used. For all γ-proteobacteria and actinobacteria competitions, 9:1 ratios (DSS-3:competitor) were used. For these experiments, log relative competitive indexes (log RCI) were calculated for each coincubation. A positive log RCI indicates that DSS-3 had a competitive advantage after coincubation, while a negative log RCI indicates that the competitor strain had an advantage. A log RCI of zero indicates that neither strain has a competitive advantage. After 24 hours, all coincubations of competitor strains with DSS-3 Kn, had significantly higher log RCI values compared to that of coincubations with the non-killer DSS-3 mutant GCS64 (Fig. 7), suggesting the GBL cluster is required for DSS-3 to outcompete these isolates in coculture. The one exception was *Microbacterium* sp. RAM275, which when coincubated with either DSS-3 Kn or GCS64 had log RCI values near zero, suggesting the DSS-3 killing mechanism does not convey a competitive advantage against this Actinobacterium under these conditions.

**Figure 7.**
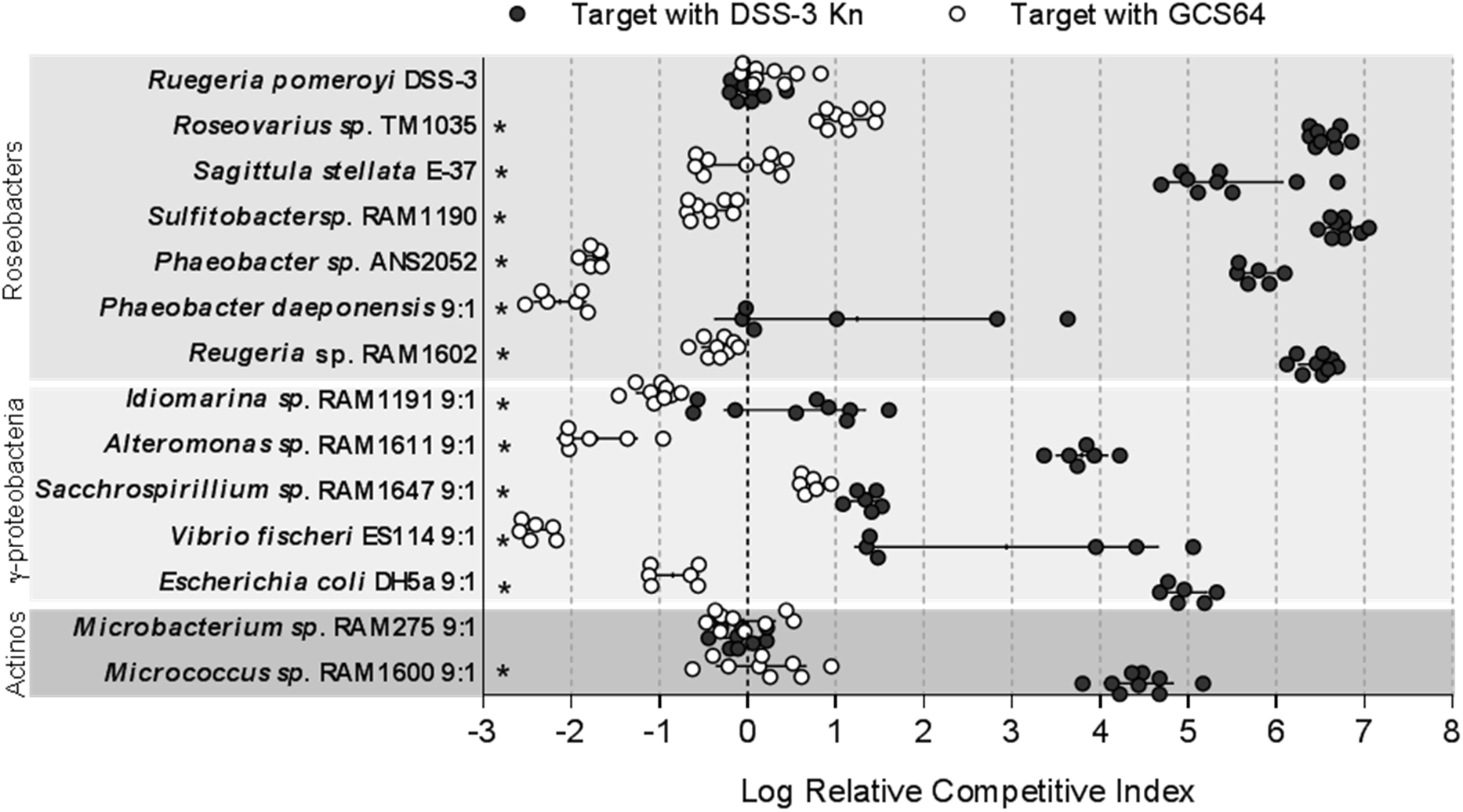
Putative GBL gene cluster is required to maintain DSS-3’s competitive advantage against phylogenetically diverse bacteria. Log relative competitive index (log RCI) for cocultures of chosen competitors with either DSS-3 Kn (black) or the *afsA* mutant GCS64 (white) after 24 hours of coincubation on surfaces. LogRCI was calculated as follows: log RCI= log((DSS-3 CFU_24HR_/Competitor CFU_24HR_)/(DSS-3 CFU_0HR_/Competitor CFU_0HR_)). A positive log RCI value corresponds to DSS-3 having a competitive advantage, and a negative Log RCI corresponds to the competitor strain having a competitive advantage. Asterisks denote a *P*<0.05 using a student’s t-test comparing the log RCI values of competitions with DSS-3 Kn or with *afsA* mutant GCS64. Error bars indicate standard deviation.

## Discussion

Based on the data presented here, we propose the following model for how the GBL gene cluster may enable DSS-3 to eliminate competitors. DSS-3 grows together in the presence of competitor bacteria until a cell density threshold is reached, at which point the antimicrobial can eliminate phylogenetically-diverse competing bacteria, significantly reducing their population sizes and enabling DSS-3 to dominate the niche space.

Both killing and immunity functions require genes encoded in the GBL gene cluster. Given that the killing phenotype is density-dependent and requires GBL biosynthesis genes, we hypothesize that a GBL-like molecule may be the antimicrobial and a high cell density is needed to achieve cytotoxic levels sufficient to kill competitor cells, or GBL may act as a signaling molecule that combines with one or more regulators to activate production of an unknown, diffusible antimicrobial whose synthesis is regulated in a density- and GBL-dependent manner. Future work should focus on identifying the antimicrobial molecule, the type of GBL produced, and its specific role in mediating interbacterial killing.

Because competition can impact microbial community structure and function, it is critical to identify the ecologically-relevant habitats and conditions that support and restrict DSS-3’s killing activity. The density dependent requirement of this killing mechanism limits the environments and micro-habitats where such a competitive mechanism may be advantageous. In our study, DSS-3 needed to achieve a cell density of ~10^8^ CFU/ml to kill a competitor. However, the cell density threshold required for killing in the marine environment may be different because environmental viscosity and cellular metabolism can influence fluid flow of the local environment, which may promote or prevent the accumulation of signaling and/or antimicrobial molecules (53, 54). For example, the phycosphere or organic particles are habitats with low diffusibility and high nutrients (55), and therefore may support growth of DSS-3 microcolonies and allow local concentrations of these molecules to activate killing. DSS-3 has already been found to establish a mutualistic relationship with the diatom *Thalassiosira pseudonana* in coculture (56): DSS-3 provides the limiting micronutrient B_12_ to *T. pseudonana*, and the diatom in turn produces the sulfur-carbon metabolite C-3 sulfonate 2,3-dihydroxypropane-1-sulfonate (DHSP) (56). This mutualistic exchange of resources between DSS-3 and *T. pseudonana* suggests that DSS-3 may indeed have the capacity to colonize certain phytoplankton. If DSS-3 does colonize the phycosphere, DSS-3 may be able to use the diffusible antimicrobial described here to kill competitors, significantly impacting the structure and function of the phytoplankton microbiome and ecophysiology.

We found several lines of evidence that suggest the GBL synthesis gene cluster in DSS-3 may have been a recent acquisition event. Specifically, we found that 1) the GBL-synthesis gene cluster is located on a megaplasmid, 2) homologs are absent in other *Ruegeria* species but present in disparate individuals within various families of α-proteobacteria, γ-proteobacteria, and actinobacteria, and 3) organisms encoding AfsA domain proteins were mostly Actinobacteria and γ-proteobacteria. It is also notable that the putative GBL gene cluster identified here has previously only been described in actinobacteria as a mechanism to regulate the production of secondary metabolites, including antimicrobials (46). Therefore, the discovery of a putative GBL synthesis gene cluster outside of Actinobacteria suggests an expanded role for GBL signaling that has not previously been considered.

In summary, this work demonstrates that *R. pomeroyi* DSS-3 encodes a broadly effective, diffusible antimicrobial that can alter the abundance of co-occurring marine bacteria, providing a mechanism by which this organism can significantly alter the structure and function of marine microhabitats. Future work is needed to identify the antimicrobial molecule as well as the ecological niches where it is employed.

## Acknowledgements

The authors would like to thank Peggy Cotter for providing plasmids; Mary Ann Moran for providing roseobacter strains; Stephanie Smith, Lauren Speare, Laura Fisch, and Daniel Efird for technical assistance; Brett Froelich for assistance in the field; Carol Arnosti for the opportunity to participate in two Atlantic cruises supported by NSF1332881 from which isolates were obtained; and the assistance of the captain, crew, and scientific parties of the R/V Endeavor cruises EN556 and EN584. GCS was supported by a University of North Carolina Graduate Merit Assistantship and a Junior Faculty Development Award to ANS.

